# Neurotrophin-3 from the dentate gyrus supports postsynaptic sites of mossy fiber-CA3 synapses and contextual memory

**DOI:** 10.1101/2023.07.16.549236

**Authors:** Ji-Wei Tan, Haifei Xu, Guey-Ying Liao, Juan Ji An, Baoji Xu

## Abstract

At the center of the hippocampal tri-synaptic loop are synapses formed between mossy fiber (MF) terminals from granule cells in the dentate gyrus (DG) and proximal dendrites of CA3 pyramidal neurons. However, the molecular mechanism regulating the development and function of these synapses is poorly understood. In this study, we showed that neurotrophin-3 (NT3) was expressed in nearly all mature granule cells but not CA3 cells. We selectively deleted the NT3-encoding *Ntf3* gene in the DG during the 1^st^ two postnatal weeks to generate a *Ntf3* conditional knockout (Ntf3-cKO). Ntf3-cKO mice had normal hippocampal cytoarchitecture but displayed elevated anxiety level and impairments in contextual memory, spatial reference memory and nest building. As MF-CA3 synapses are essential for encoding of contextual memory, we examined synaptic transmission at these synapses using ex vivo electrophysiological recordings. We found that Ntf3-cKO mice showed impaired basal synaptic transmission due to deficits in excitatory postsynaptic currents mediated by AMPA receptors but normal presynaptic function and intrinsic excitability of CA3 pyramidal neurons. Consistent with this selective postsynaptic deficit, Ntf3-cKO mice had fewer and smaller thorny excrescences on proximal apical dendrites of CA3 neurons and lower GluR1 levels in the stratum lucidum area where MF-CA3 synapses reside but normal MF terminals, compared with control mice. Thus, our study indicates that NT3 expressed in the dentate gyrus is crucial for the postsynaptic structure and function of MF-CA3 synapses and hippocampal-dependent memory.

## Introduction

The hippocampus is critical for encoding, storing, and retrieving of contextual memory [1, 2]. Information from the entorhinal cortex (EC) is processed in the hippocampus through several circuits, including the well-known tri-synaptic loop: EC → dentate gyrus (DG) → CA3 → CA1, which provides inputs back to the EC. At the center of this loop are synapses formed between mossy fiber (MF) terminals from DG granule cells and proximal dendrites of CA3 pyramidal neurons. CA3 pyramidal neurons also receive perforant path inputs directly from the EC and recurrent inputs from other CA3 pyramidal neurons onto their distal dendrites. MF-CA3 synapses are essential for encoding of contextual memory [3–5]; however, the molecular mechanism regulating the development and function of these synapses is poorly understood.

MF-CA3 synapses have unique properties regarding their development, structure, and function. First, the DG is one of two brain regions with active adult neurogenesis [6, 7] so that granule cells and MF-CA3 synapses are constantly generated throughout lifespan. Second, the synapses have large MF terminals with multiple neurotransmitter release sites that contact large, complex, and multi-headed dendritic spines called thorny excrescences (TEs) [8, 9]. Lastly, long-term potentiation (LTP) at MF-CA3 synapses is independent of NMDA receptors and is entirely expressed presynaptically, which is in contrast with LTP at many other synapses such as CA1 synapses [10, 11]. Brain-derived neurotrophic factor (BDNF), one of neurotrophins that also include nerve growth factor (NGF), neurotrophin-3 (NT3), and neurotrophin-4/5 (NT4/5), is known to regulate the development and function of several synapses in response to neuronal activity [12, 13]. However, the role of neurotrophins in the development and function of MF-CA3 is not clear, although BDNF has been reported to regulate maturation of adult-born granule cells [14, 15]. Despite NT3 is expressed in the DG [16, 17] and its receptor TrkC is expressed in both the DG and CA3 region [18], the role of NT3 in the development and function of MF-CA3 synapses has not been determined yet.

In this study, we employed the Pomc-Cre transgene [19] to selectively abolish NT3 expression in the DG to generate conditional knockout mice. We found that mutant mice displayed elevated anxiety and deficits in contextual memory and nest building. Interestingly, the NT3 abolishment reduced the size of TEs and excitatory postsynaptic currents (EPSCs) mediated by the α-amino-3-hydroxy-5-methyl-4-isoxazolepropionic acid receptor (AMPAR) at MF-CA3 synapses without affecting the function of presynaptic sites. These results reveal a critical role for NT3 in the structure and function of postsynaptic sites and contextual memory.

## Materials and Methods

### Animals

The following mouse strains were obtained from the Jackson Laboratory: *Ntf3^lox/lox^* (stock No: 003541), Pomc-Cre (stock No: 010714), Ai9 (stock No: 007909), Thy1-GFP (stock No: 00778), and wild-type C57BL/6J (stock No: 000664). The *Ntf3^LacZ/+^* mouse strain was obtained from MMRRC (ID No: 000191-UCD). Mice were group housed and maintained at 22 °C on a 12-h/12-h light/dark cycle with *ad libitum* access to food and water. All animal procedures used in this study were approved by the UF Scripps Institutional Animal Care and Use Committee.

### In situ hybridization

Mice at 6 weeks of age were used for in situ hybridization, which was performed as described previously [20]. In brief, mouse brains were dissected and frozen immediately in an isopentane-dry ice bath. All in situ hybridization was performed on cryostat coronal sections at 10 µm. To generate antisense riboprobes, mouse cDNA sequence for *Ntf3* (GenBank accession number NM_008742, nucleotides 214–990) was amplified by PCR and cloned into the pBluscript II KS (−) plasmid (Stratagene, La Jolla, CA). Antisense RNAs were synthesized from linearized plasmids using T3 RNA polymerase (New England Biolabs, Ipswich, MA), [α-35S]-CTP, and [α-35S]-UTP (PerkinElmer, Waltham, MA). X-ray films were scanned into digital images at 1200 dpi.

### Immunohistochemistry and Nissl staining

Mice were deeply anaesthetized with avertin and transcardially perfused with phosphate buffered saline (PBS), followed by 4% paraformaldehyde (PFA) in PBS. Brains were post-fixed in 4% PFA for 6 h, and then cryoprotected in 30% sucrose in PBS until sectioning. Coronal brain sections (40 μm in thickness) were obtained using a sliding microtome (Leica SM2000R). Brain sections were rinsed once with Tris buffered saline (TBS; 10 mM Tris-HCl, 150 mM NaCl, pH7.5), incubated with blocking buffer (0.4% Triton X-100, 1% bovine serum albumin, and 10% goat serum or horse serum in TBS) for 1 h at room temperature, and then incubated with primary antibodies diluted in the blocking buffer overnight at room temperature. The following primary antibodies were used: chicken anti-β-galactosidase (1:3,000; Abcam #ab9361), rabbit anti-calbindin (1:400; Cell Signaling Technology #13176S), goat anti-doublecortin (1:200; Santa Cruz Biotechnology #SC-8066), rabbit anti-GluR1 (1:250; MilliporeSigma #ab1504). For GluR1 staining, sections were digested in 0.2 M HCl containing 0.5 mg/ml pepsin (MilliporeSigma, #10108057001) for 10 min at 37 °C, and then transferred to the blocking buffer as described [21]. After three washes in TBS, sections were incubated with appropriate fluorescent secondary antibodies (1:500; Jackson ImmunoResearch) for 1 h at room temperature. Sections were washed three times in TBS, mounted onto slides, and coverslipped with a mounting medium containing 4′,6-diamidino-2-phenylindole (DAPI; Vector Laboratories). Fluorescent images were captured using a Nikon C2+ confocal microscope. Quantitative analysis was performed by using Image J (NIH software) or manually by an individual blind to genotype. Nissl staining was performed by submerging mounted sections in cresyl violet for 20 min before dehydration as described [15].

### Stereology

We used Stereo Investigator software (MBF Bioscience, Inc., Williston, VT) to estimate cell density of Nissl-stained brain samples as described [22]. Measurements were performed on every fourth stained coronal sections throughout the hippocampus. Both hemispheres of a given brain were analyzed separately. The number of neurons in the DG, CA3, and CA1 was estimated using a fractionator sampling method. For each stereological probe, neurons were counted within a counting frame of 40 µm x 40 µm (a sampling site), and 4–10 sampling sites were randomly picked by the software within the outlined area. The counts were then extrapolated to estimate the cell density in the DG, CA3 and CA1. For all probe runs the coefficient of error (CE Scheaffer) was <1.

### Imaging of Thy1-GFP brains

Mice at 8-12 weeks old were deeply anaesthetized with avertin and transcardially perfused with phosphate buffered saline (PBS), followed by 4% PFA in PBS. Brains were removed, post-fixed in 4% PFA at 4 °C for 6 h, and then cryoprotected in 30% sucrose in PBS at 4 °C until sectioning. Coronal brain sections (60 μm thick) were cut on a sliding microtome (Leica SM2000R) and collected in PBS. Sections were washed in PBS, mounted onto slides, and coverslipped with a mounting medium containing 4′,6-diamidino-2-phenylindole (DAPI; Vector Laboratories).

Images of GFP fluorescence were acquired with a Nikon C2+ confocal microscope. Images were obtained with identical acquisition settings regarding exposure time, detector gain, and amplifier offset. For quantification, GFP fluorescence intensity in the MF was normalized to the intensity in the DG cell layer [23]. For MF terminals and dendritic spines, images were taken from the stratum lucidum (SL) and stratum radiatum (SR) layers of CA3. Images at 1024 × 1024 pixels were acquired as a z-stack (0.5 µm step size) using 100x with 1.0x zoom for MF terminals or 60x with 2.0x zoom for dendritic spines. The size (area) of each MF bouton was measured using Image J. The number of MF boutons was counted manually by an experimenter blind to the genotype. The number of MF boutons was divided by the SL area of each image for comparison. The number of proximal TEs and distal dendritic spines was counted manually. The number of dendritic spines was divided by the dendrite length. The size (area) of each TEs was measured using Image J.

### Stereotaxic injection of viruses for sparse cell labeling

We obtained pAAV-Ef1a-FLPo (Addgene plasmid #55637, a gift from Karl Deisseroth) and produced AAV5-Ef1a-FLPo through the UNC Vector Core. AAV2-Ef1a-fDIO-mCherry were purchased from Addgene (Addgene viral prep #114471-AAV2). To sparsely label neurons for morphological analysis, diluted AAV5-Ef1a-FLPo (1:500) was mixed with AAV2-Ef1a-fDIO-mCherry at equal volume. The virus mixture (200 nl) was injected bilaterally into the CA3 of 8-12-week-old mice at a rate of 25 nl per min using Microsyringe Pump (World Precision Instruments) as described previously [24]. Stereotaxic coordinates from the bregma and skull used wwere as follows: AP: −1.50 mm, ML: ± 2.40 mm, and DV: −2.35mm. Three weeks after virus injection, mice were perfused with PBS and then 4% paraformaldehyde in PBS. Brains were removed, post-fixed in 4% PFA at 4 °C for 6 h, and then cryoprotected in 30% sucrose in PBS at 4 °C until sectioning. Coronal brain sections at 60-μm thickness were cut using a sliding microtome (Leica SM2000R) and collected in PBS. Sections were washed in PBS, mounted onto slides, and coverslipped with a mounting medium containing 4′,6-diamidino-2-phenylindole (DAPI; Vector Laboratories).

### 2D TE morphology measurements

TE morphology measurements were conducted by using Nikon-elements Analysis software. Proximal dendritic segments were chosen from the somas to the end of TE locations (within stratum lucidum, SL). The area of TEs was calculated by measuring the total area (TEs + dendrites) within Z-stack images then subtracting the dendritic area from the total. For each cell, TE density was calculated as the total TE area divided by total dendritic length measured [5].

### Electrophysiology

Mice were decapitated under isoflurane anesthesia. The brain was rapidly removed and placed in ice-cold (4 °C) artificial cerebrospinal fluid (aCSF) containing (in mM): 124 NaCl, 3 KCl, 26 NaHCO_3_, 1.25 NaH_2_PO_4_, 1 MgSO_4_, 2 CaCl_2_, and 10 D-glucose, equilibrated with 95% O_2_ and 5% CO_2_. Transverse hippocampal slices (300 μm thick) were obtained using a vibratome (Leica VT 1200s, Germany) and then transferred to oxygenated aCSF at 32 °C for recovery.

Mice at 2-3 months of age were used for field potential recordings. Slices were incubated in oxygenated aCSF at 32 °C for at least 30 min, then maintained at room temperature (23-25 °C) for another 30 min before recording. Slices were gently transferred to the recording chamber (RC-27, Warner Instruments, Hamden, CT) at room temperature. Chamber was perfused with non-circulated oxygenated aCSF at a flow rate of 2–3 ml/min. Field excitatory postsynaptic potentials (fEPSPs) were evoked by a concentric bipolar stimulating electrode (inner diameter: 25 μm; outer diameter: 125 μm, FHC Inc., Bowdoin, ME) connected to a constant current stimulus isolator (SYS-A365R, WPI, Sarasota, FL). Recordings were performed with low resistance (1–3 MΩ) glass pipettes (ID: 0.68 mm, OD: 1.2 mm, WPI, Sarasota, FL) filled with aCSF under current-clamp mode with I=0 configuration. To obtain fEPSPs at hippocampal MF-CA3 synapses, both stimulating and recording electrodes were placed in the SL of the CA3 sub-region as described [23]. For input–output measurement, fEPSP slope was recorded by increasing the stimulation intensity (0.1-ms pulse width) from 0 to 100 μA in a 10-μA increment. For paired-pulse ratio (PPR) and long-term potentiation (LTP), stimulation intensities were adjusted to give 50% of the maximal fEPSP slopes. PPRs were obtained by delivering paired stimulating pulses to the synapses with inter-pulse interval from 50 ms to 200 ms in a 50-ms increment. The value of PPRs was calculated as the slope of the second synaptic response was divided by the slope of the first synaptic response. LTP of the MF-CA3 synapses was evoked by high-frequency stimulation (HFS) consisting of four trains (1 s at 100 Hz for each train) applied at 20-s intervals after 20-min stable baseline recording at 0.033-Hz test stimulation. LTP was recorded for 1 h after stimuli applied. Data of LTP was expressed as averages of fEPSP slope for every 2 min of recordings, and slopes in the last 10 min of recordings were averaged per animal.

Mice at P21-P28 were used for whole-cell recordings. CA3 pyramidal neurons were visually identified in slices using an infrared-differential interference contrast microscope (Scientifica, UK). Whole-cell patch-clamp recordings were performed using borosilicate glass pipettes of 3–5 MΩ pulled with a micropipette puller (P-1000; Sutter Instrument, Novato, CA). For voltage-clamp recording, pipettes were filled with internal solution containing (in mM): 115 CsMeSO_3_, 20 CsCl, 10 Hepes, 0.6 EGTA, 4 Mg-ATP, 0.3 Na_3_-GTP, 1 QX-314, 2.5 MgCl_2_, and 10 Na_2_-Phosphocreatine (pH 7.3 with CsOH, osmolarity 285 mM). Miniature excitatory postsynaptic currents (mEPSCs) were recorded by holding neurons at −70 mV without series resistance and liquid junction compensation. Neurons with Ra>25 MΩ were excluded. mEPSCs were recorded in the presence of 1 μM tetrodotoxin (TTX, American Radiolabeled Chemicals, Inc., #ARCD 0640) and 100 μM picrotoxin (PTX, Sigma-Aldrich, # P1675). For the amplitude and frequency of mEPSCs measurements, the detection threshold was set at 4 pA and 100 events were sampled per neuron. Evoked AMPAR- and N-methyl-D-aspartate NMDA receptor (NMDAR)-mediated EPSCs were recorded by holding neurons at −70 mV and +40 mV, respectively, in the presence of 100 μM picrotoxin. Stimulating electrodes were placed in the SL layer of the CA3 subregion. The input-output curves of AMPAR- and NMDAR-mediated EPSCs were recorded by increasing the stimulation intensity (0.1-ms pulse width) from 20 μA to 45 μA in 5-μA increments. The AMPAR/NMDAR ratio was calculated by measuring EPSC amplitudes at the same stimulus intensity from the same cell. AMPAR-mediated EPSCs were measured as the peak responses following the stimulus, and NMDAR-mediated EPSCs were measured as the mean current over a 5-ms window, 50 ms after the stimulus as described [25]. Current-clamp recording was used to test excitability of the CA3 pyramidal neurons. The pipettes for current-clamp recording were filled with internal solution containing (in mM): 130 potassium gluconate, 6 NaCl, 1 MgCl_2_, 20 Hepes, 0.2 EGTA, 2 Mg-ATP, 0.3 Na_3_-GTP (pH 7.3 with KOH, osmolarity 285 mM). Membrane capacitance, series resistance, and holding current were determined from the Clampex 10.6 readout during recording. Resting membrane potential (RMP) was obtained immediately by switching to current-clamp after whole-cell mode was established. Input resistance was measured by the slope of the linear fit of the I-V curve from −200 to 0 pA current injection for 1 s in 50-pA increment. The sag ratio was calculated as the ratio of steady-state voltage change to the maximum voltage change caused by hyperpolarizing current injection (−200 pA for 1s) as described [26]. Neuronal excitability was evaluated by counting the number of spikes evoked in response to a series of 1-s depolarizing current injection steps (ranging from 0 to 250 pA in 50-pA increments with a 30-s intertrial interval). Action potential threshold was read out from each single action potential waveform [27].

The experimenter was blind to genotype. Signals were acquired with Multiclamp 700B and Digidata 1550A (Molecular Devices, San Jose, CA). Data were low-pass filtered at 2.9 KHz and sampled at 10 kHz. fEPSPs, mEPSCs, I-V curves and action potentials were analyzed with Clampfit 10.6 software.

### Behavioral tests

Mice at 2–4 months of age were tested during the dark (active) phase of a 12-/12-h reversed light– dark cycle. Mice were transferred to a holding room in the behavioral testing area, at least 1 h before the behavioral assays. Each apparatus was cleaned with 1% Micro-90 between every trial. Offline automatic behavioral analysis was performed using the Ethovision XT video tracking system (Noldus, Netherlands). Different tests were spaced by at least 3 days with less stressed tests being conducted first.

#### Open-field test

The mouse was gently placed in a corner of an open-field box (43.8 cm × 43.8 cm × 32.8 cm) facing the corner and monitored voluntary activity for 30 min. Total distance moved, velocity and center zone duration were recorded automatically.

#### Elevated plus-maze test

The apparatus consisted of two open arms (35 cm × 6 cm), two closed arms (35 cm × 6 cm with 20 cm walls) and a central zone (6.1 cm × 6.1 cm). The plus-shaped maze was elevated 60 cm from the floor. The mice were placed on the central zone facing one open arm and allowed to explore freely for 5 minutes. The time spent in open arms and entries into open arms were recorded automatically.

#### Light–dark box test

A black plastic insert with lid was placed into the box (60 cm x 40.0 cm x 25.0 cm), thus the apparatus was divided into two equal zones (a brightly lit compartment and a dark one). The black insert had a door that allowed animals to shuttle freely. Mice were placed in the bright chamber facing the wall and allowed to explore the two chambers for 5 min. Latency to enter the dark chamber, duration of time in the light chamber and number of crossings between chambers were automatically recorded.

#### Contextual fear conditioning

The mouse was placed into a Phenotyper chamber (29.2 cm x 29 cm x 30.5 cm, Noldus) equipped with an electrified floor and speaker enclosed in a ventilated and sound-attenuated box. Training consisted of a 150 s baseline followed by three 0.75 mA foot shocks. 24 hours after conditioning, the mice were tested for contextual fear memory by reexposing them to the training chamber for 5 min. Freezing behavior was automatically recorded.

Pain threshold to footshock was tested as described [28]. Mice were placed into the conditioning chamber, and 3 minutes later they received footshocks of increasing amplitude spaced 30 seconds apart. Starting with 0.05 mA, footshock was increased in 0.05-mA increments until four response thresholds were reached: flinching (hind/fore paws briefly raise off the bars), move, vocalizing, and jumping.

#### Morris water maze

Spatial learning and memory was assessed in the MWM using a video-tracking system (EthoVision XT, Noldus), and was performed as described [29]. Mice were subjected to swim for 60 s to find the hidden escape platform (10 cm in diameter) in a pool (120 cm in diameter, water temperature was held at 23 °C) surrounded by distal visual cues. If a mouse failed to find the platform within a 60-s trial, it was guided to the platform by the experimenter, and remained there for 15 s. Testing was conducted in three phases: visible platform training (day 0), acquisition training (days 1-7, four trials with different starting points per day) and probe trial (day 8, without the platform). Escape latency, time spent in the target quadrant, latency to the platform location, platform area crossing numbers and swimming speed were recorded.

#### Forced swimming test

A mouse was placed in a transparent plastic cylinder (13 cm in diameter by 24 cm in height) filled with 10-cm depth of water at room temperature. Its behavior was recorded by a video-tracking system (EthoVision XT, Noldus) for a total of 6 min. The total duration when the mouse was immobile was recorded.

#### Tail suspension test

A mouse was suspended from a hook by the tail with an adhesive tape in a three-walled compartment. The tail base area was covered with a transparent plastic tail guard to prevent tail climbing behavior. Mouse behavior was recorded by a video-tracking system (EthoVision XT, Noldus) for a total of 6 min. The total duration when the mouse was immobile was measured.

#### Three-chamber sociability and social novelty preference test

The three-chamber test for sociability and response to social novelty was performed as previously described [25, 29]. Each chamber was 20 cm x 40 cm x 25 cm in size, with rectangular openings (5 cm x 8 cm) on the central dividing walls made of clear acrylic. During habituation, a test mouse was placed in the central chamber and allowed to freely explore the empty apparatus for 5 min. Immediately after habituation, a stimulus mouse (same-sex, similar age, and unfamiliar mouse) was introduced into a wire cage (7 cm in diameter and 15 cm in height with bars spaced 1 cm apart) located in one of the two side chambers. An identical empty wire cage was placed in the other side chamber. The test mouse was allowed to explore the arena for 10 min. After the first 10-min trial, each mouse was tested in a second 10-min trial to evaluate social preference for a different strange mouse. Another unfamiliar mouse was introduced to the previously empty cage. The test mouse had a choice between the first, already-investigated mouse (familiar), and the novel unfamiliar mouse (stranger). Time spent in sniffing the stimulus mouse (or empty wire cage) was recorded by Ethovision XT video tracking system. In addition to analyzing investigation time, we also calculated the social preference index and social discrimination index, which represent the differences between time spent in investigating the stimulus mouse vs. object and familiar mouse vs. stranger mouse, divided by total time spent in investigating both targets, respectively.

#### Nest building test

Nest building behavior was assessed in regular mouse housing cage after a mouse was individually housed for a week. Approximately 1 hour before the dark cycle, mice were transferred to test cages with nesting materials (Nestlet, 2.6 g, 5 cm x 5 cm and 5 mm thick compressed cotton). Nest construction was scored at the beginning of the light cycle with a 5-point system: 1, nestlet not noticeably touched; 2, nestlet partially torn up (50–90% remaining intact); 3, mostly shredded nestlet but often no identifiable nest site; 4, an identifiable, but flat nest; 5, a (near) perfect nest with clear nest crater as described [30]. Quantification of the normalized weight of the remaining unshredded nesting material was performed as an objective measure of nest building.

### Statistical analysis

Results are expressed as the mean ± SEM unless indicated otherwise. Significance was set at P < 0.05 for all statistical measures. All data were evaluated using a 2-tailed Student’s t test, 2-tailed Mann-Whitney test or 2-way ANOVA with post hoc analyses where applicable. All graphs and statistics were made using Graph Pad Prism 9.0 software. All figures were created using Adobe Illustrator.

## Results

### *Ntf3* gene expression and deletion in the DG

We used adult *Ntf3^LacZ/+^* knockin mice [31] for examination of NT3 expression in the DG. Immunohistochemistry revealed that nearly all mature granule cells marked by calbindin (94.2 ± 0.8%, n=3 mice) expressed NT3 as indicated by β-galactosidase (β-gal) (Fig. 1A). Similarly, 94.2% NT3-expressing DG cells were mature granule cells (Fig. 1A, B) and 2.3% of them immature granule cells expressing DCX (Fig. 1C, D). Many immature granule cells were negative for β-gal immunoreactivity (Fig. 1C). Thus, NT3 is expressed in nearly all mature granule cells and a fraction of immature granule cells.

**Fig. 1.**
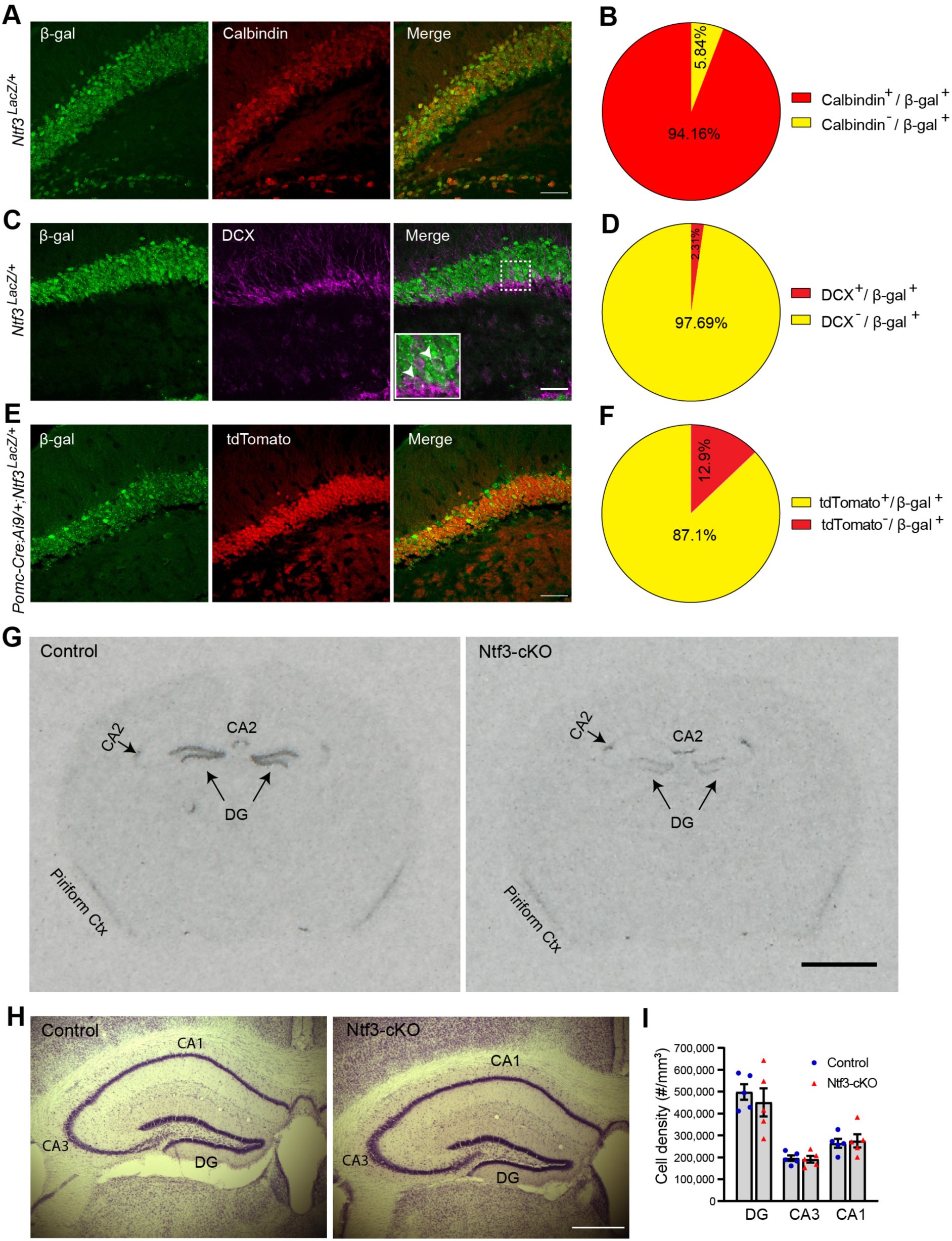
Deletion of the *Ntf3* gene in DG granule cells. (**A** and **B**) Colocalization of β-galactosidase (β-gal) with calbindin in the DG of *Ntf3^LacZ/+^*mice (n = 3 adult mice). Scale bar, 50 µm. (**C** and **D**) Colocalization of β-gal with DCX in the DG of *Ntf3^LacZ/+^* mice (n = 3 adult mice). Scale bar, 50 µm. (**E** and **F**) Colocalization of β-gal with tdTomato in Pomc-Cre;Ai9;*Ntf3^LacZ/+^* mice (n = 3 adult mice). Scale bar, 50 μm. (**G**) Representative images of *Ntf3* mRNA in situ hybridization in brain sections of adult *Ntf3^lox/lox^* (control) and Pomc-Cre*;Ntf3^lox/lox^* (Ntf3-cKO) mice. Scale bar, 2 mm. (**H**) Nissl staining of brain sections showing normal hippocampal cytoarchitecture in Ntf3-cKO mice. Scale bar, 500 µm. (**I**) Cell density in the hippocampal DG, CA3, and CA1 subregions in control and Ntf3-cKO mice at 3 months of age. n = 5 mice per group; two-way ANOVA, p = 0.486. Error bars represent SEM.

To determine when DG granule cells start to express NT3, we conducted calbindin and β-gal immunohistochemistry on brain sections from *Ntf3^LacZ/+^* mice at P0, P7, P14, and P21. We did not detect β-gal immunoreactivity in the DG at P0 (Supplementary Fig. S1A); however, the level of β-gal immunoreactivity in the DG at P7 was comparable to those at P14 and P21, with approximately 90% of β-gal-immunoreactive cells being mature granule cells (Supplementary Fig. S1B-G). Thus, DG granule cells start to express NT3 during the first postnatal week, and the expression quickly reaches the peak.

To delete the *Ntf3* gene in the DG, we used a Pomc-Cre transgene in which Cre expression begins in the first postnatal week and is mostly restricted to DG granule cells [19, 32]. To further examine the timing and efficiency of recombination mediated by Pomc-Cre, we crossed Pomc-Cre mice to Ai9 mice, which are a Cre reporter mouse strain designed to have a loxP-flanked STOP cassette preventing transcription of a CAG promoter-driven tdTomato-coding gene until Cre-mediated recombination between the two loxP site occurs [33]. In agreement with previous studies [19, 32], tdTomato expression was restricted to the DG in Pomc-Cre/+;Ai9/+ mice (Supplementary Fig. S2A). In newborn Pomc-Cre/+;Ai9/+ pups, the DG area contained tdTomato-expressing cells but very few calbindin-expressing mature granule cells (Supplementary Fig. S2B), suggesting that Pomc-Cre is expressed in immature granule cells or their precursors. We observed that 16%, 80%, and 89% mature granule cells expressed tdTomato in 1-week-old, 2-week-old, and adult Pomc-Cre;Ai9 mice, respectively (Supplementary Fig. S2C-H), indicating deletion of a loxP-flanked gene in mature granule cells is almost complete by 2 weeks of age. In adult Pomc-Cre;Ai9;*Ntf3^LacZ/+^* mice, 87% of β-gal-expressing DG cells also expressed tdTomato (Fig. 1E, F), indicating that Pomc-Cre can be used to efficiently abolish NT3 expression in the DG.

We generated control (*Ntf3^lox/lox^*) and DG-specific Ntf3 conditional knockout Pomc-Cre/+; *Ntf3^lox/lox^* (Ntf3-cKO) mice by crossing female *Ntf3^lox/lox^* mice to male Pomc-Cre/+;*Ntf3^lox/+^*mice, because Pomc-Cre has been found to be expressed female germline cells [19]. In situ hybridization (ISH) revealed that levels of NT3 mRNA in Ntf3-cKO mice were greatly reduced in the DG but not in the CA2 compared with *Ntf3^lox/lox^* control (Ctrl) mice (Fig. 1G). Pomc-Cre is also expressed in a few other brain regions such as the arcuate nucleus of the hypothalamus and lateral habenular nucleus [19]; however, these brain regions do not express NT3 (Allen Brain Atlas ISH Data). Thus, NT3 expression in Ntf3-cKO mice is selectively abolished in the DG during the first 2 postnatal weeks. Stereological analysis found that *Ntf3* deletion in the DG did not alter the cytoarchitecture and cell density of the hippocampus (Fig. 1H, I).

### Memory deficits in Ntf3-cKO mice

Male Ntf3-cKO mice moved normally in open field tests (Fig. 2A), indicating that *Ntf3* deletion in the DG does not impair motor function. These mice had anxiety-like behaviors, as they spent less time in the open arm and entered the open arm fewer times in elevated plus maze tests (Fig. 2B, C) and spent less time in the light chamber in light-dark box tests (Fig. 2D, E). We run contextual fear conditioning tests (Fig. 2F) and water maze tests (Fig. 2J) to determine if *Ntf3* deletion in the DG affects contextual memory and spatial reference memory. Ntf3-cKO mice spent a lower percentage of time in freezing than control mice during the 2^nd^ and 3^rd^ footshock trials (Fig. 2G) and when returning to the training chamber 24h after conditioning (Fig. 2H), and their sensitivity to footshock was normal (Fig. 2I), indicating that Ntf3-cKO mice have deficits in contextual learning and memory. In the visible platform water maze test, Ntf3-cKO mice performed normally (Fig. 2K), indicating that vision and swimming ability are not affected in mutant mice. In the hidden platform of water maze test, control and cKO mice had similar escape latency during 7-day training (Fig. 2L). However, in day-8 probe trial Ntf3-cKO mice showed a trend of spending less time in the target quadrant, took longer time to reach the platform location for the 1^st^ time, and crossed the platform location fewer times than control mice (Fig. 2M-O). These results show that *Ntf3* deletion in the DG impairs spatial reference memory. Furthermore, we noticed that male Ntf3-cKO had deficits in nest building (Fig. 2P-R) although they had normal body temperature (Fig. 2S). Lesions of the hippocampus, medial preoptic area, and septum have been found to impair nesting [34–37]. Thus, the nesting deficit in male Ntf3-cKO mice could be related to hippocampal dysfunction.

**Fig. 2.**
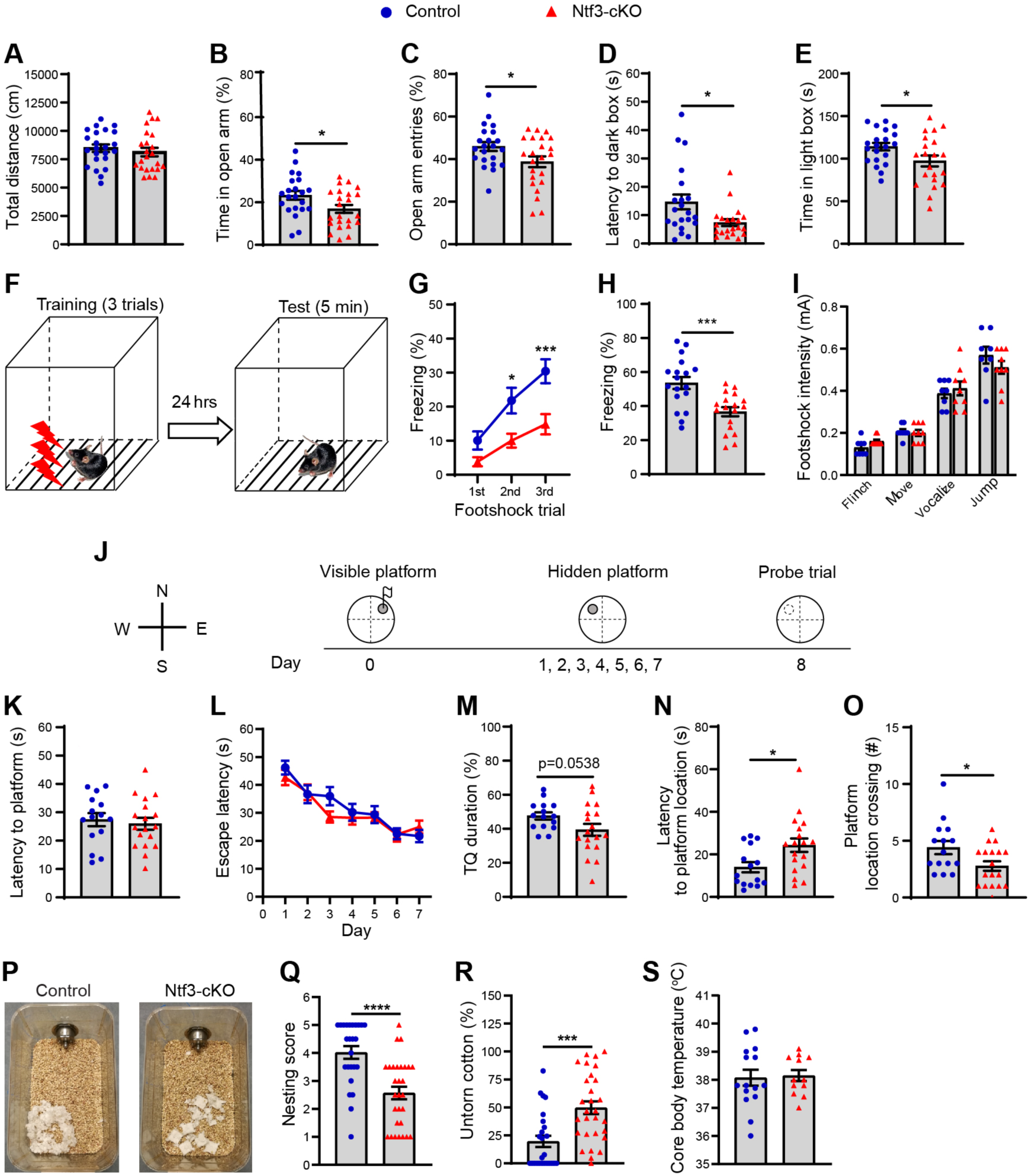
Elevated anxiety level and impaired hippocampus-dependent cognitive function in male Ntf3-cKO mice. (**A**) Total travel distance of mice in open field tests. n = 22 mice per genotype; Two-tailed unpaired t test, p=0.705. (**B** and **C**) Time spent in the open arm and frequency of entry into the open arm in elevated plus-maze tests. n = 22 control mice and 23 Ntf3-cKO mice; Two-tailed unpaired t test, p = 0.0261 for time in the open arm and p = 0.0359 for open arm entries. (**D** and **E**) Latency to enter the dark box and time spent in the light box in light-dark box tests. n = 21 mice per genotype; Two-tailed unpaired t test, p = 0.0172 for latency to enter the dark box and p = 0.0365 for time spent in the light box. (**F** to **I**) Contextual fear conditioning test. (**F**) Schematic illustration for contextual fear conditioning test. (**G**) Percentage of time in freezing during contextual fear conditioning. n = 18 mice per genotype; Two-way ANOVA with post hoc Bonferroni’s multiple comparisons, p = 0.0019 for genotype. (**H**) Percentage of time freezing during contextual fear memory tests performed 24 hours after conditioning. n = 18 per genotype; Two-tailed unpaired t test, p = 0.0005. (**I**) Pain threshold to footshock in control and Ntf3-cKO mice. n = 8 control mice and 9 Ntf3-cKO mice; Two-way ANOVA with post hoc Bonferroni’s multiple comparisons, p = 0.879 for genotype. (**J** to **O**) Morris water maze test. (**J**) Schematic illustration of Morris water maze test. (**K**) Escape latency in visible platform water maze tests. n = 15 control mice and 18 Ntf3-cKO mice; Two-tailed unpaired t test, p = 0.64. (**L**) Escape latency in hidden platform water maze tests. n = 15 control mice and 18 Ntf3-cKO mice. Two-way ANOVA with post hoc Bonferroni’s multiple comparisons, p = 0.281 for genotype. (**M**) Percentage of time spent in in the target quadrant (TQ) during the probe trial. n = 15 control mice and 18 Ntf3-cKO mice; Two-tailed unpaired t test, p = 0.0538. (**N**) Latency to reach the platform location during the probe trial. n = 15 control mice and 18 Ntf3-cKO mice; Two-tailed unpaired t test, p = 0.0173. (**O**) Number of platform location crossing during the probe trial. n = 15 control mice and 18 Ntf3-cKO mice; Two-tailed unpaired t test, p = 0.0264. (**P** to **S**) Nest building test. (**P**) Representative nests built by control and Ntf3-cKO mice. (**Q**) Scores of nests built by control and Ntf3-cKO mice. n = 25 control mice and 29 Ntf3-cKO mice; Two-tailed Mann Whitney test, p < 0.0001. (**R**) Percentage of untorn cotton after 12-hour nest building period. n = 25 control mice and 29 Ntf3-cKO mice; Two-tailed unpaired t test, p = 0.0003. (**S**) Core body temperature. n = 15 control mice and 12 Ntf3-cKO mice; Two-tailed unpaired t test, p = 0.835. *p<0.05, **p<0.01, ***p<0.001, and ****p<0.0001. Error bars represent SEM.

Ntf3-cKO mice displayed neither depression-like behavior in forced swimming tests and tail suspension tests nor deficits in social behaviors in sociability and social novelty tests (Supplementary Fig. S3). Female Ntf3-cKO mice also displayed deficits in contextual learning and memory (Supplementary Fig. S4H-J) and nest building (Supplementary Fig. S4K-N) but did not show anxiety- or depression-related behaviors (Supplementary Fig. S4A-G).

We also examined Pomc-Cre/+ mice and wild-type (WT) littermates in contextual fear conditioning tests and nest building tests. Pomc-Cre/+ and WT mice performed comparably in these two tasks (Supplementary Fig. S5). This result rules out the possibility that the observed abnormal behaviors in Ntf3-cKO mice result from the Pomc-Cre transgene. Collectively, the behavioral tests indicate that *Ntf3* deletion in the DG leads to hippocampal dysfunction and deficits in contextual memory in mice of both sexes and elevates anxiety levels in male mice.

### Impaired postsynaptic function at MF-CA3 synapses of Nft3-cKO mice

As MF-CA3 synapses are essential for encoding of contextual memory [3–5], the deficit in contextual memory in Ntf3-cKO mice may result from dysfunction of MF-CA3 synapses. We examined the function of MF-CA3 synapses in Nft3-cKO mice at 8-12 weeks of age. We first compared field excitatory postsynaptic potentials (fEPSPs) in the SL area of the CA3 subregion evoked by stimulation at MFs in slices from control and Ntf3-cKO mice (Fig. 3A). We plotted the amplitude of the presynaptic fiber volley (input) against the slope of fEPSPs (output) over a range of stimulus strengths. The input-output curve indicates that basal synaptic transmission at MF-CA3 synapses is impaired in Ntf3-cKO mice (Fig. 3B). To determine if the synaptic deficit is due to impaired neurotransmitter release, we measured LTP (Fig. 3C) and paired-pulse facilitation (PPF; Fig. 3D) at MF-CA3 synapses. PPF is a form of short-term synaptic plasticity and reflects changes in the properties of presynaptic terminals [38], whereas LTP at MF-CA3 synapses is entirely expressed presynaptically [10]. Both forms of synaptic plasticity were normal in Ntf3-cKO mice (Fig. 3C, D), indicating that ablation of NT3 expression in the DG does not affect the function of MF terminals. In addition, *Ntf3* deletion did not affect intrinsic excitability of CA3 pyramidal neurons (Supplementary Fig. S6).

**Fig. 3.**
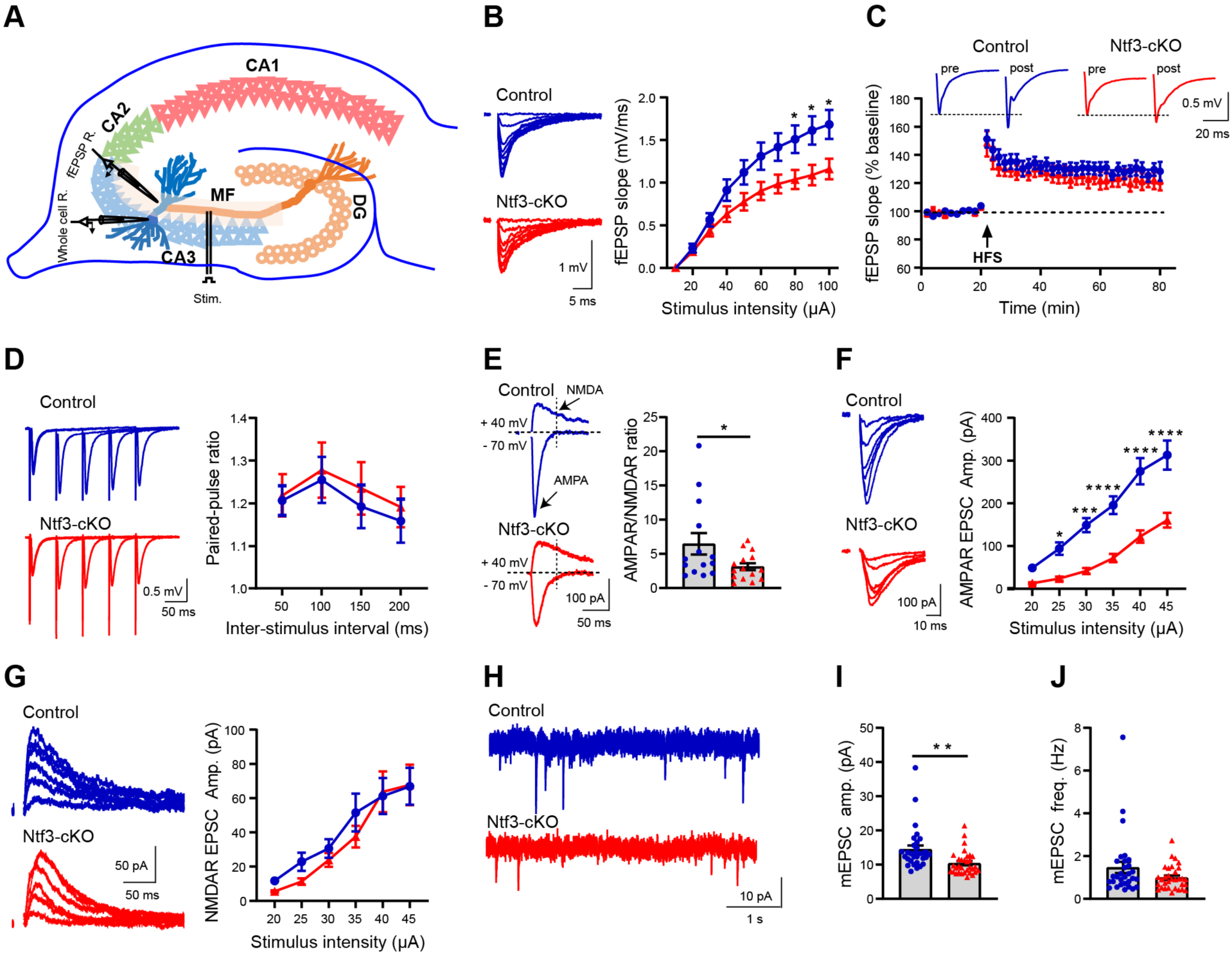
Impaired postsynaptic function at MF-CA3 synapses in Ntf3-cKO mice. (**A**) Diagram of field EPSP (fEPSP) recordings and whole-cell recordings at MF-CA3 synapses. Field recordings and whole-cell patch-clamp recordings were done in brain slices from mice at 8-12 months and 3-4 weeks old, respectively. (**B**) Input-output curves of fEPSPs, showing impaired basal synaptic transmission at MF-CA3 synapses in Ntf3-cKO mice. Representative traces are shown in the left side. n = 16 slices from 9 control mice and 15 slices from 9 Ntf3-cKO mice. Two-way ANOVA with post hoc Bonferroni’s multiple comparisons, p = 0.0381 for genotype; *p < 0.05. (**C**) LTP induced by high-frequency stimulation. Representative traces before and after stimulation are shown at the top. n = 10 slices from 7 mice of each genotype. Two-tailed unpaired t test, p = 0.349 for the last 10 minutes of LTP. (**D**) Measurement of paired-pulse facilitation at MF-CA3 synapses. Representative traces are shown in the left side. n = 12 slices from 6 control mice and 14 slices from 8 Ntf3-cKO mice. Two-way ANOVA with post hoc Bonferroni’s multiple comparisons, p=0.468 for genotype. (**E**) AMPAR-EPSCs to NMDAR-EPSCs ratio. Whole-cell recordings with holding potentials at −70 mV and 40 mV were used to measure AMPAR-EPSCs and NMDAR-EPSCs, respectively. n = 14 cells from 7 control mice and 15 cells from 7 Ntf3-cKO mice. Two-tailed unpaired t test, p=0.0466; *p < 0.05. (**F**) Input-output curve of AMPAR-EPSCs recorded at −70 mV. Representative traces are shown in the left side. n = 25 cells from 9 control mice and 22 cells from 11 Ntf3-cKO mice. Two-way ANOVA with post hoc Bonferroni’s multiple comparisons, p<0.0001 for genotype; *p<0.05, ***p<0.001, and ****p<0.0001. (**G**) Input-output curve of NMDAR-EPSCs recorded at 40 mV. Representative traces are shown in the left side. n = 17 cells from 7 control mice and 18 cells from 8 Ntf3-cKO mice. Two-way ANOVA with post hoc Bonferroni’s multiple comparisons, p<0.51 for genotype. (**H**-**J**) Recordings of mEPSCs. Representative traces (**H**), amplitude of mEPSCs (**I**), and frequency of mEPSCs (**J**) are shown. n = 30 cells from 8 control mice and 32 cells from 9 Ntf3-cKO mice. Two-tailed unpaired t test; p = 0.0027 for amplitude and p = 0.0843 for frequency; **p<0.01. Error bars represent SEM.

We then investigated if the deficit at MF-CA3 synapses results from postsynaptic impairments. We performed whole-cell patch-clamp recordings in CA3 neurons to measure AMPAR-EPSCs (holding potentials = −70 mV) and NMDAR-EPSCs (holding potentials = 40 mV) in the presence of 100 μM picrotoxin on brain slices from mice at 3-4 weeks of age. The ratio of AMPAR-EPSCs over NMDAR-EPSCs was lower in Ntf3-cKO mice than in control mice (Fig. 3E). Input-output curves for AMPAR-EPSCs (Fig. 3F) and NMDAR-EPSCs (Fig. 3G) revealed that the reduction in the AMPAR-EPSC/NMDAR-EPSC ratio was due to diminished AMPAR-EPSCs in Ntf3-cKO mice. These results indicate that NT3 deficiency leads to impairments in the function of AMPA receptors at MF-CA3 synapses.

CA3 pyramidal neurons in Ntf3-cKO mice at 3-4 weeks age exhibited reduced amplitude but normal frequency of miniature EPSCs (mEPSCs), which were measured in the presence of 1 μM tetrodotoxin and 100 μM picrotoxin, compared with those in control mice (Fig. 3H-J). As the amplitude and frequency of mEPSCs are, respectively, reflective of postsynaptic and presynaptic functions, this result further supports the conclusion that NT3 deficiency impairs postsynaptic function at MF-CA3 synapses without affecting presynaptic function. Because impairments in both AMPAR-EPSCs and mEPSCs in Ntf3-cKO mice were detectable by 3-4 weeks of age, NT3 must regulate the development of the postsynaptic sites of MF-CA3 synapses. This regulation appears to be specific to glutamatergic synapses, as the amplitude and frequency of miniature inhibitory postsynaptic currents (mIPSCs) were normal in CA3 pyramidal neurons of Ntf3-cKO mice (Supplementary Fig. S7).

### Reduced number and size of TEs in Ntf3-cKO mice

To understand the structural basis of dysfunction at MF-CA synapses, we employed the Thy1-GFP transgene (M transgenic line) [39] to examine MF terminals and spines on apical dendrites of CA3 pyramidal neurons. Thy1-GFP is expressed in many granule cells and labels MFs (Fig. 4A). GFP fluorescence intensity of MFs in the CA3 area was comparable between control and Ntf3-cKO mice (Fig. 4A, B), indicating that *Ntf3* deletion in the DG does not affect the number of MFs. Furthermore, the size and density of MF terminals in the CA3 SL area were also normal in Ntf3-cKO mice (Fig. 4C-F). Conversely, Ntf3-cKO mice had fewer and smaller TEs, which form synapses with MF terminals, on the proximal dendrites of CA3 pyramidal neurons (Fig. 4G-I) but normal density of dendritic spines on the distal dendrites of CA3 pyramidal neurons which form synapses with axons from the EC and recurrent collaterals from the CA3 (Fig. 4J, K). Because very few CA3 pyramidal neurons express Thy1-GFP (Fig. 4A) and GFP-labeled MFs interfere with visualization of GFP-labeled TEs, we further examined TEs using viruses to sparsely label CA3 pyramidal neurons. A mix of FLPo-dependent AAV2-Ef1a-fDIO-mCherry and diluted FLPo-expressing AAV5-Ef1a-FLPo was bilaterally injected into the CA3 of control and Ntf3-cKO mice at about 2 months of age (Fig. 4L). The amount of TEs on proximal dendrites of CA3 pyramidal neurons was significantly reduced (Fig. 4M, N). These results show that NT3 expressed in granule cells is released at MF terminals and act on CA3 pyramidal neurons to regulate the development and/or maintenance of TEs.

**Fig. 4.**
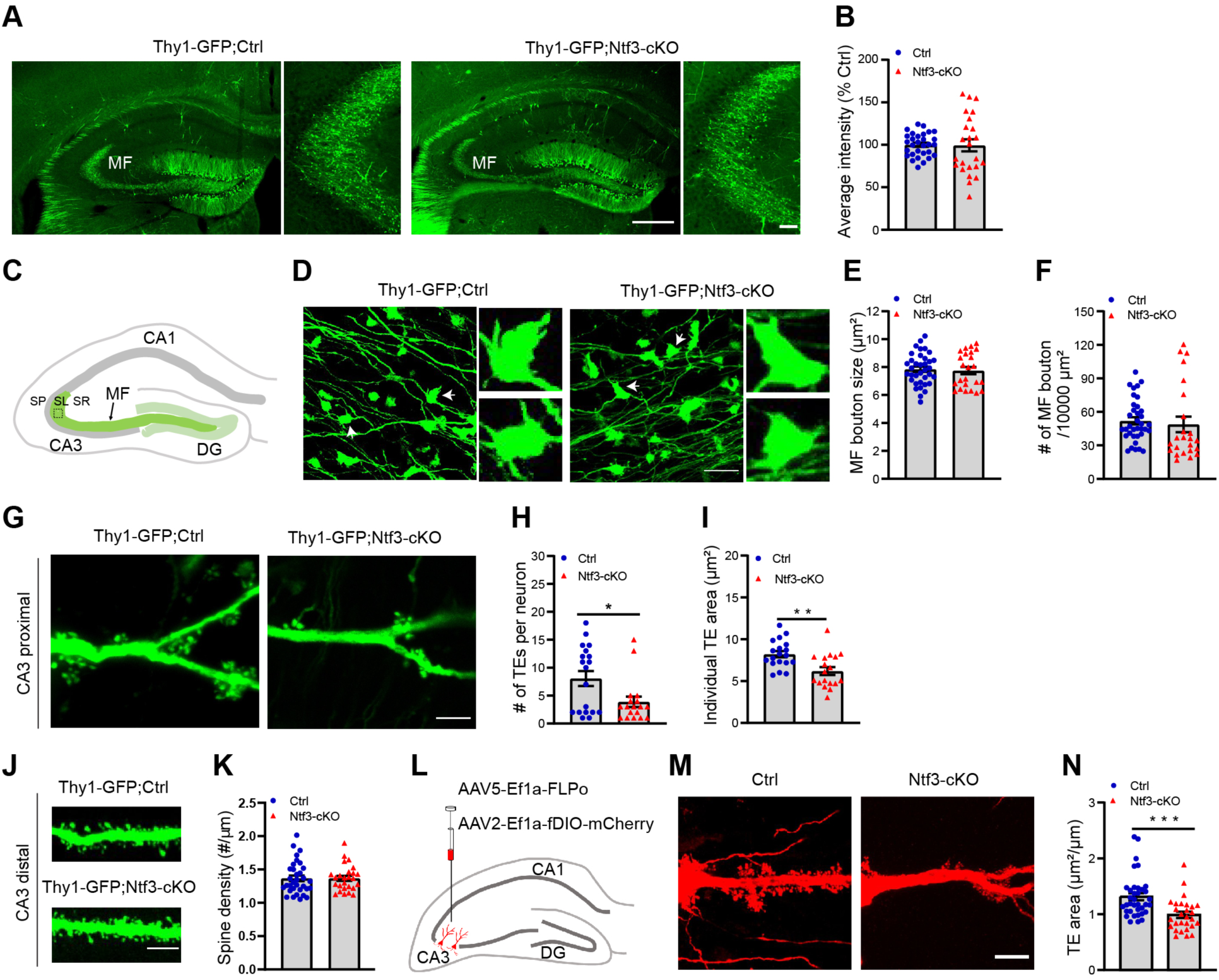
Deletion of the *Ntf3* gene in DG granule cells reduces the number and size of thorny excrescences. (**A)** Representative images of Thy1-GFP-labeled granule cells and MF. Scale bars, 500 μm for larger images and 50 μm for smaller images. (**B**) Quantification of GFP fluorescence in the MF of the CA3 subregion. n = 30 fields from 5 Thy1-GFP;control mice and 24 fields from 4 Thy1-GFP;Ntf3-cKO mice. Two-tailed unpaired t test, p = 0.918. (**C** to **F**) Size and density of MF terminals. (**C**) Schematic diagram indicating the pictured area in (**D**), (**E**), and (**F**). Images were taken from the SL area of the CA3 subregion (boxed). (**D**) Representative images of MF boutons. MF boutons denoted by arrowheads are magnified. Scale bar, 10 μm. (**E** and **F**) Normal size and density of MF boutons in Ntf3-cKO mice. n = 36 fields from 6 Thy1-GFP;control mice and 24 fields from 4 Thy1-GFP;Ntf3-cKO mice. Two-tailed unpaired t test, p = 0.745 for bouton size and 0.642 for bouton density. (**G**) Representative images of Thy1-GFP-labeled proximal dendrites of CA3 pyramidal neurons, showing thorny excrescences (TEs). (**G**) Representative images of proximal apical dendrites of CA3 pyramidal neurons. Scale bar, 10 μm. (**H** and **I**) Density and size of TEs. n = 19 neurons from 5 Thy1-GFP;control mice and 18 neurons from 5 Thy1-GFP;Ntf3-cKO mice. Two-tailed unpaired t test, p=0.0163 for density of TEs and p=0.00197 for size of TEs; *p<0.05 and **p<0.01. (**J**) Representative images of Thy1-GFP-labeled distal apical dendrites of CA3 pyramidal neurons. Scale bar, 10 μm. (**K**) Spine density on distal apical dendrites of CA3 pyramidal neurons. n = 26 dendrite segments from 7 Thy1-GFP;control mice and 33 dendrite segments from 9 Thy1-GFP;Ntf3-cKO mice. Two-tailed unpaired t test, p = 0.994. (**L**) Schematic illustration of AAV5-Ef1a-FLPo-WPRE and AAV2-Ef1a-fDIO-mCherry stereotaxically injected into the CA3 regions to label pyramidal neurons. (**M**) Representative images of proximal apical dendrites of mCherry-labeled CA3 pyramidal neurons. Scale bar, 10 µm. (**N**) Ntf3-cKO mice showed significantly smaller TE area than their controls. n=36 neurons from 5 Ctrl mice and 28 neurons from 5 Ntf3-cKO mice. Two-tailed unpaired t test, p<0.001. Error bars represent SEM.

### Reduced levels of GluA1 at MF-CA3 synapses of Ntf3-cKO mice

We performed GluA1 immunohistochemistry to determine whether diminished AMPAR-EPSCs result from a reduced amount of AMPA receptors at MF-CA3 synapses which reside in the SL area (Fig. 5A). Quantitative analysis of confocal images using ImageJ software founds that the size and number of GluA1^+^ puncta in Ntf3-cKO mice were reduced in the SL area (Fig. 5B-D) but not in the SR area (Fig. 5E-G) which receive perforant path inputs directly from the EC and recurrent inputs from other CA3 pyramidal neurons onto their distal dendrites. As levels of GluA1 in the SR area was not altered, it is likely that GluA1 gene expression is normal in CA3 pyramidal neurons of Ntf3-cKO mice. Therefore, these results suggest that NT3 from granule cells regulates trafficking of AMPA receptors into postsynaptic sites in complex and multi-headed TEs.

**Fig. 5.**
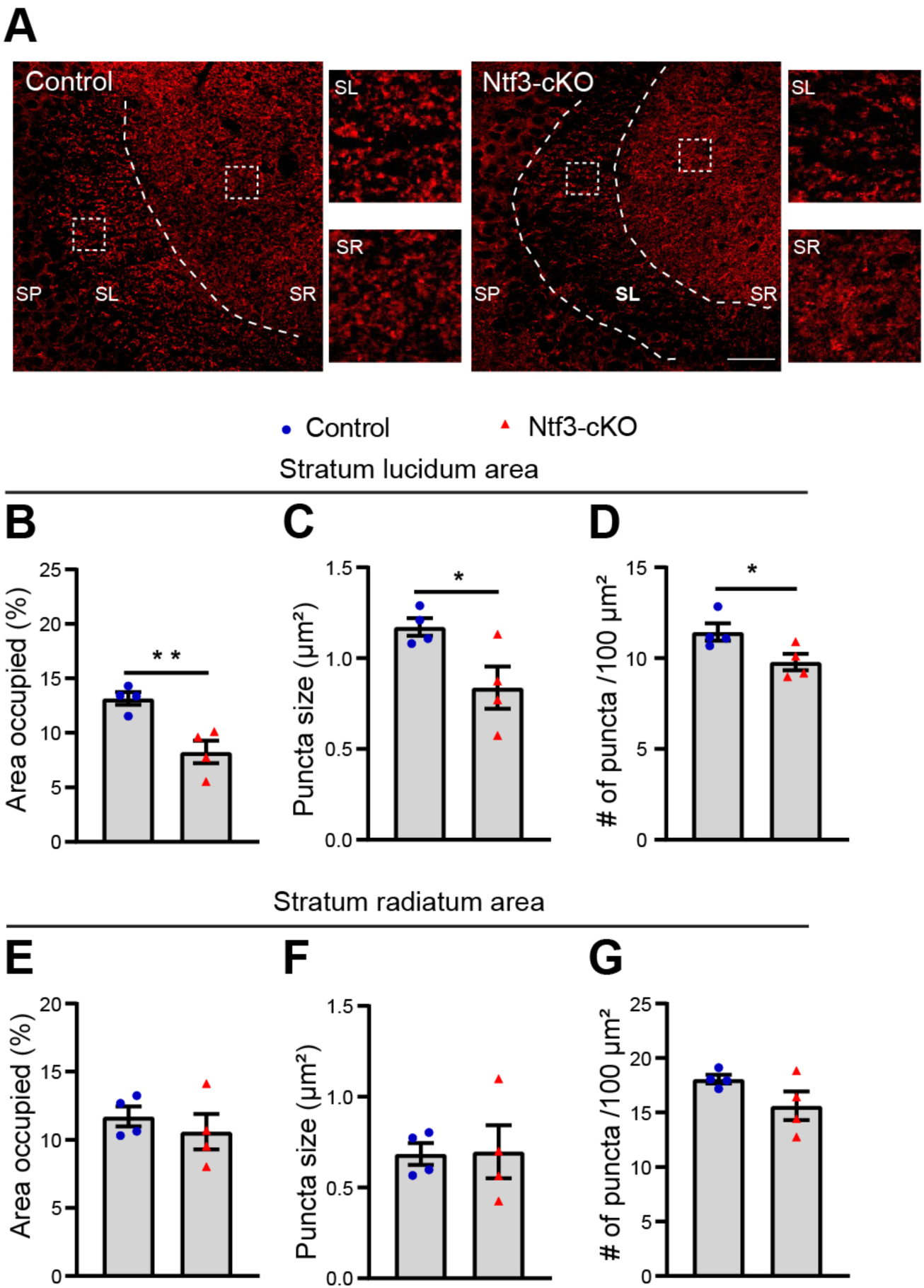
GluA1 levels are reduced in the SL area but not the SR area of the CA3 subregion in Ntf3-cKO mice. (**A**) Representative confocal images of GluA1 immunoreactivity in the CA3 subregion. Scale bar, 50 µm. (**B**-**D**) Area occupied by GluA1 immunoreactivity, size of GluA1 immunoreactive puncta, and density of GluA1 immunoreactive puncta in the SL area. n = 4 mice per genotype. Two-tailed unpaired t test, *p < 0.05 and **p < 0.01. (**E**-**G**) Area occupied by GluA1 immunoreactivity, size of GluA1 immunoreactive puncta, and density of GluA1 immunoreactive puncta in the SR area. n = 4 mice per genotype. Two-tailed unpaired t test, p = 0.481 for area occupied, 0.939 for puncta size, and 0.123 for density of puncta. Error bars represent SEM.

## Discussion

Studies on the role of neurotrophins in the CNS have almost completely focused on BDNF. NT3 is expressed the cingulate cortex, piriform cortex, hippocampal CA2, substantia nigra, ventral tegmental area, and cerebellum in addition to the DG [16, 17, 40]. However, very little is known about the role of NT3 in the brain. NT3 has been shown to be essential for normal cerebellar foliation [41] and dendritic morphogenesis of cerebellar Purkinje cells [42]. One study investigated the role of NT3 in hippocampal plasticity by deleting the *Ntf3* gene in all neurons during embryogenesis with Nestin-Cre and found deficits in LTP at synapses between the lateral perforant path and granule cells [43]. The study examined neither the mechanism underlying the LTP deficit nor the role of NT3 in the development, function, and structure of CNS synapses. Here, we found that ablation of NT3 expression in the DG granule cells caused postsynaptic abnormalities in structure and function at MF-CA3 synapses, including reduced size and number of TEs and diminished AMPAR-mediated EPSCs.

The postsynaptic abnormalities at MF-CA3 synapses of Ntf3-cKO mice likely result from developmental defects. We found that DG granule cells start to express NT3 after birth and the expression quickly reaches the peak by one week of age. This expression timing is coincident with formation and maturation of MF-CA3 synapses. In addition to its role as a receptor tyrosine kinase in activating several signaling cascades such as mitogen-activated protein kinase (MAPK), phosphoinositide-3-kinase (PI3K) and phospholipase C gamma (PLCψ) pathways [44], TrkC has been shown to be a synaptic organizer. Trans interaction of TrkC in postsynaptic cells with axonal protein tyrosine phosphatase PTPα generates bidirectional adhesion and formation of glutamatergic synapses, where TrkC mediates clustering of postsynaptic molecules in dendrites whereas PTPα causes differentiation of presynaptic sites [45]. Although this TrkC activity is independent of the tyrosine kinase domain and all TrkC isoforms have this activity [45], NT3 enhances the synaptic organizing activity of the TrkC-PTPα complex [46]. Therefore, it is possible that DG NT3 promotes development of MF-CA3 synapses by enhancing the activity of the TrkC-PTPα complex. This mechanism would predict that deletion of the *Ntf3* gene in the DG should also impair presynaptic function of MF-CA3 synapses. However, we observed normal paired-pulse facilitation, mEPSC frequency, and LTP at MF-CA3 synapses in Ntf3-cKO mice, all of which are dependent on presynaptic function [10, 38]. Moreover, PTPα is not expressed in DG granule cells and CA3 neurons [47]. Thus, it is more likely that NT3 is released from MF terminals, binds to TrkC at proximal dendrites of CA3 pyramidal neurons, and activates the three canonical TrkC signaling cascades to promote development of MF-CA3 postsynaptic sites.

How does NT3 promote the development of postsynaptic sites at MF-CA3 synapses? TrkC activation in proximal dendrites of CA3 pyramidal neurons can promote formation and growth of dendritic spines by stimulating protein synthesis either locally in dendrites or globally in cell bodies. As we observed that dendritic spines at distal dendrites of CA3 pyramidal neurons are normal in Ntf3-cKO mice, it is unlikely that TrkC activation in dendrites signals back to increase global gene expression in cell bodies of CA3 pyramidal neurons. Conversely, BDNF has been shown to induce local translation of mRNAs encoding proteins that are important for synaptic function and dendritic spine development, such as CamKIIalpha, NMDAR subunits, and cytoskeleton-associated protein Homer2, in the synaptodendritic compartment via the PI3K-Akt-mammalian target of rapamycin (mTOR) pathway [48]. NT3 can regulate development and function of MF-CA3 synapses through the same mechanism because binding of NT3 to TrkC also activates the PI3K-Akt-mTOR pathway. Local protein synthesis in dendrites does not explain all synaptic phenotypes we observed in Ntf3-cKO mice. As local translation of mRNAs encoding both AMPAR subunits and NMDAR subunits occurs in dendrites [48, 49], reduced local protein synthesis should impair both AMPAR-EPSCs and NMDAR-EPSCs in Ntf3-cKO mice; however, we found that *Ntf3* deletion in the DG impairs AMPAR-EPSCs but not NMDAR-EPSCs at MF-CA3 synapses. During LPT induction at Schaffer collateral-CA1 synapses, Ca^2+^ influx through the NMDAR induces incorporation of intracellular AMPAR into synapses, leading to strengthening of synaptic transmission [50]. At MF-CA3 synapses where LTP is independent of NMDA receptors [10], we speculate that NT3-TrkC signaling can modulate strength of synaptic transmission by regulating trafficking of the AMPAR. The structure of MF-CA3 synapses is highly plastic [51] and is readily modified by learning, stress, and aging [52–57]. It would be intriguing to determine whether stress and aging alter the function of MF-CA3 synapses and contextual learning and memory by regulating *Ntf3* gene expression in future studies.

## Author Contributions

J.W.T. and B.X. designed research; J.W.T., H.X., G.Y.L., and J.J.A. performed experiments and analyzed data; and B.X. and J.W.T. wrote the paper.

## Data Availability Statement

Original data generated and analyzed during this study are included in this published article.

## Acknowledgments

We thank Alicia Brantley for assistance in running behavioral tests. This work was supported by the grants from the National Institutes of Health to BX (R01 DK103335, R01 DK105954, and R01 MH125187).

## Conflicts of Interest

The authors declared that they have no competing financial interests.

## Notes

### Competing Interest Statement

The authors have declared no competing interest.

