## Supplemental Figures for "Neurotrophin-3 from the dentate gyrus supports postsynaptic sites of mossy fiber-CA3 synapses and contextual memory"

### Supplementary Information

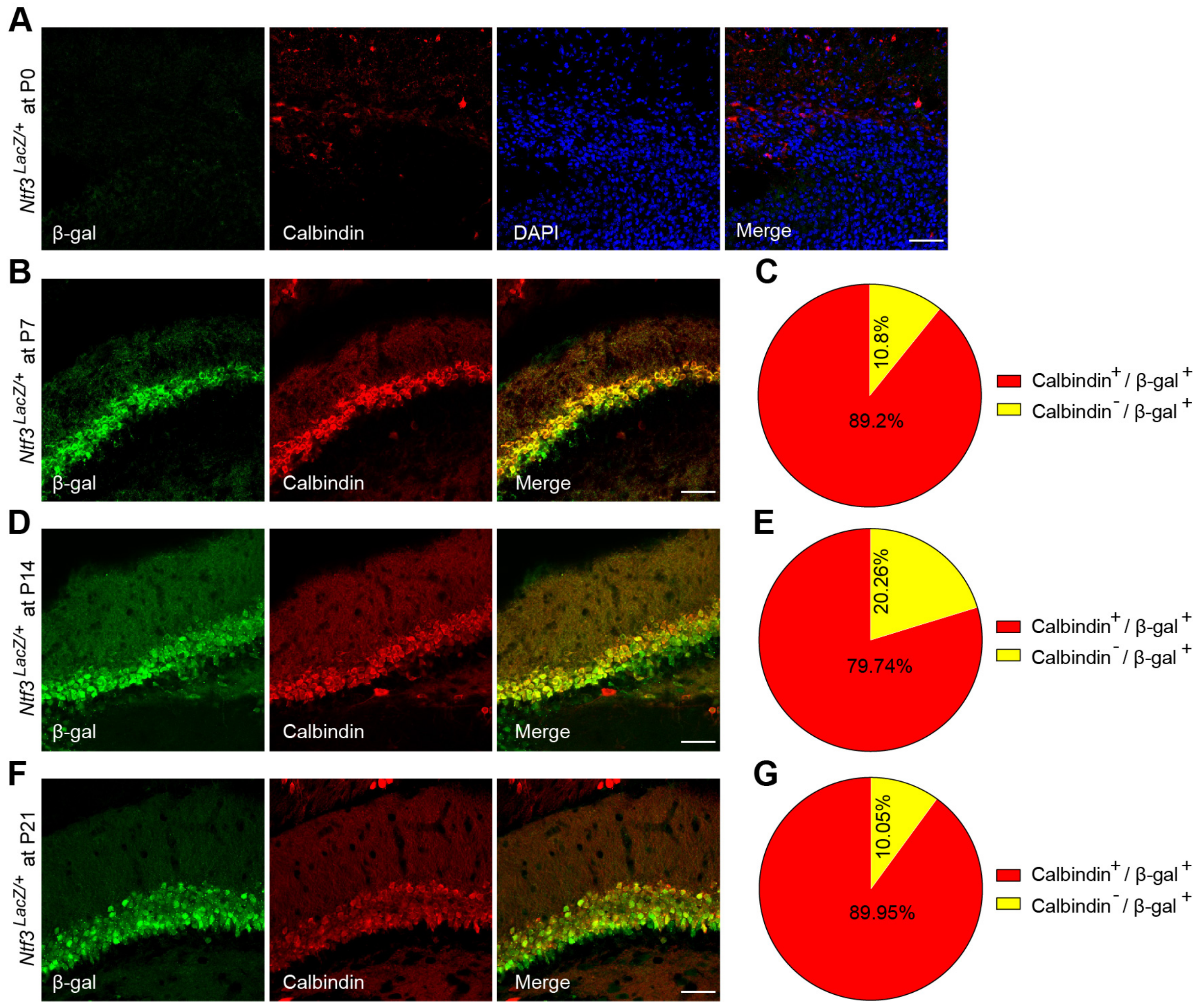


**Fig. S1 Expression of NT3 in the dentate gyrus.** NT3 expression was examined using double immunofluorescent immunohistochemistry for β-galactosidase and calbindin in brain sections from *Ntf3*$^{LacZ/+}$ mice at P0 (A), P7 (B, C), P14 (D, E), and P21 (F, G). n=2 mice for each age. Scale bars, 50 μm.

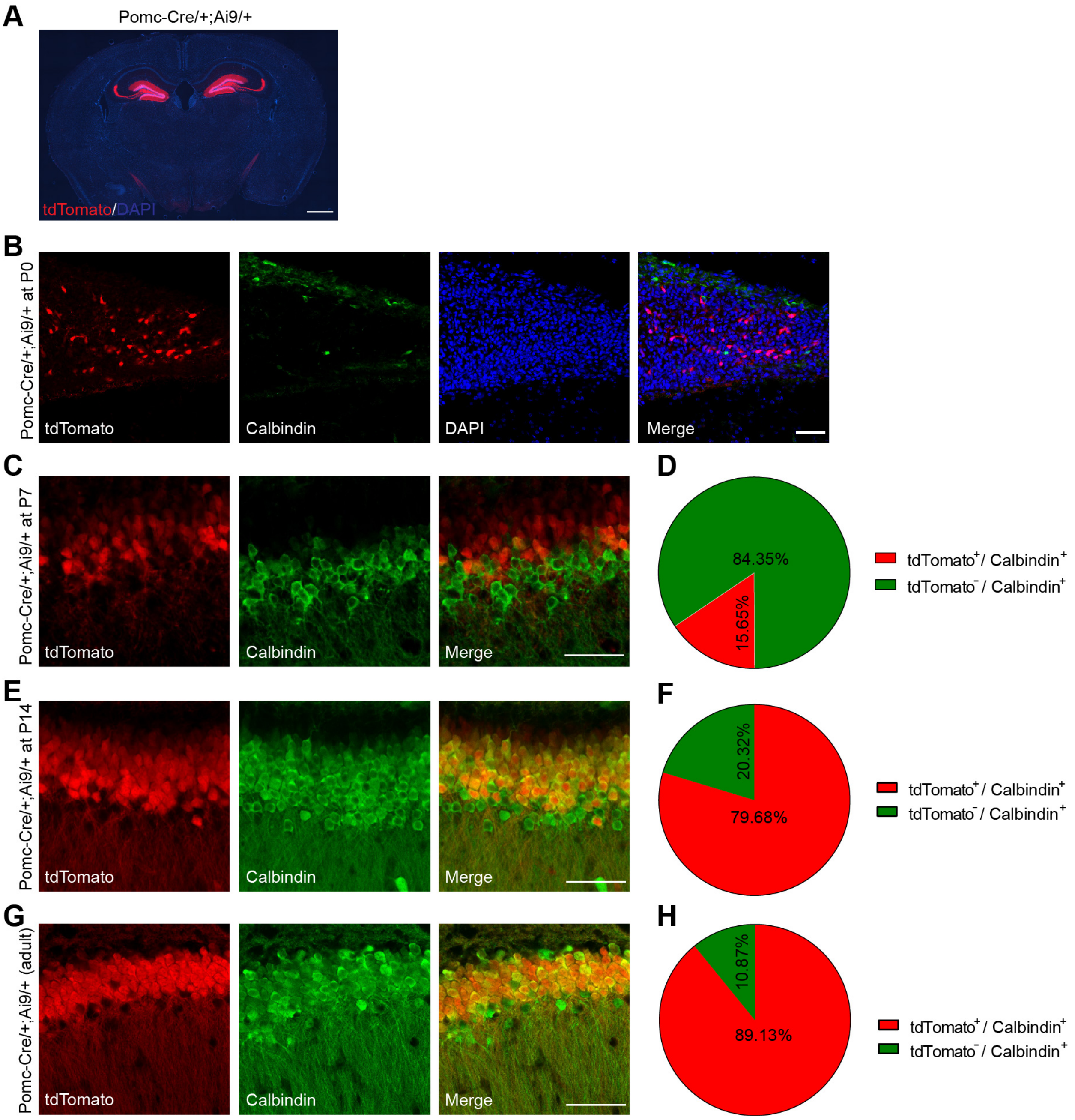


**Fig. S2 Gene deletion mediated by Pomc-Cre in hippocampal granule cells.** (**A**) tdTomato marks Cre-mediated recombination pattern in the brain of a Pomc-Cre/+;Ai9/+ mouse. Scale, 1000 μm. (**B, C, E,** and **G**) Colocalization of calbindin immunofluorescence with tdTomato in hippocampal granule cells of *Pomc-Cre/+;Ai9/+* mice at P0, P7, P14, and adult. Scale bars, 50 μm. (**D, F,** and **H**) Quantification of colocalization between calbindin immunofluorescence and tdTomato in hippocampal granule cells of Pomc-Cre/+;Ai9/+ mice at P7, P14, and adult. n = 2, 3, 2, and 5 mice at P0, P7, P14, and adult, respectively.

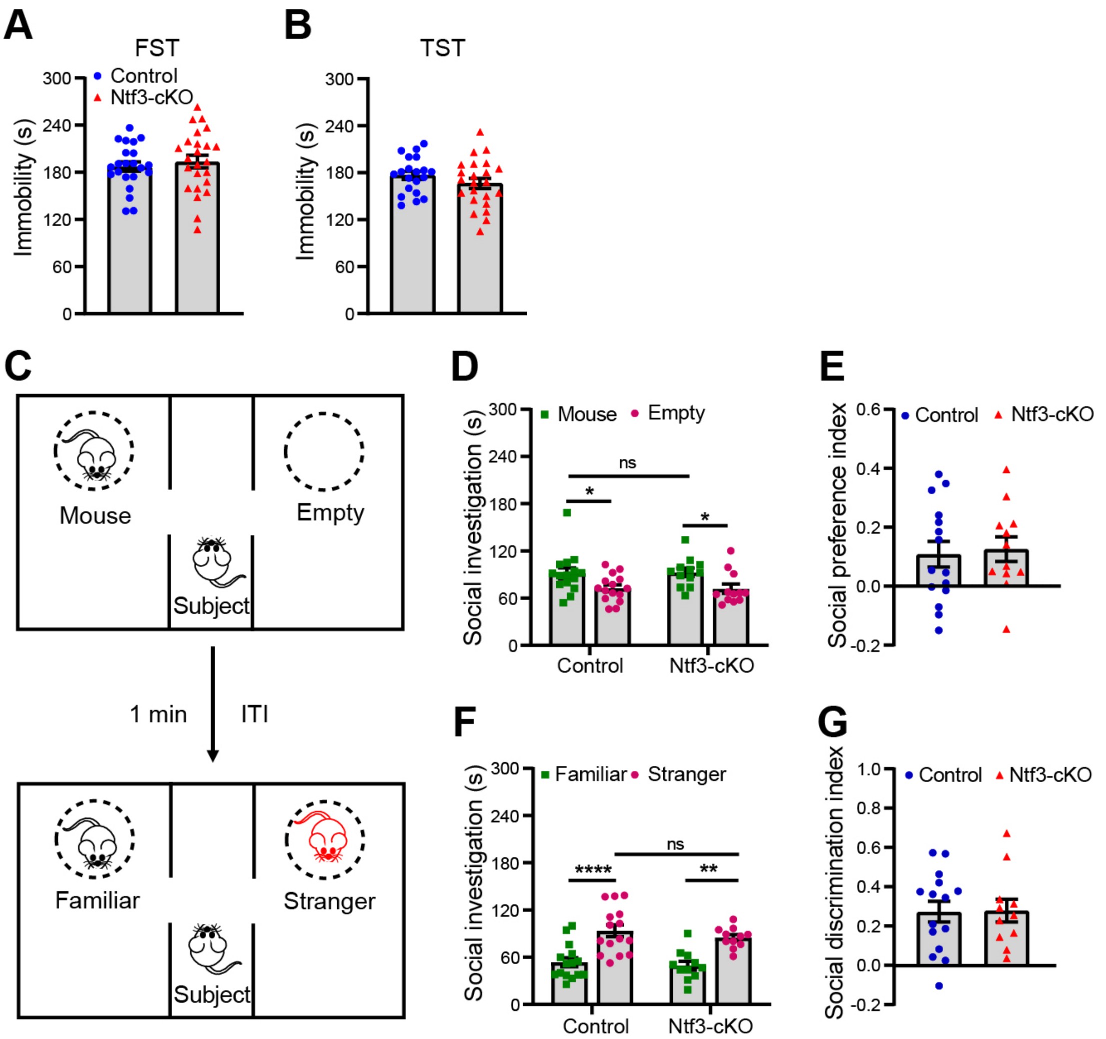


**Fig. S3 Deletion of the *Ntf3* gene in the DG doesn't affect depressive-like behaviors and social behaviors in male mice.** (**A** and **B**) Ntf3-cKO mice and control mice showed comparable immobility time in FST and TST. n = 20-24 mice per genotype. Two-tailed unpaired t test, p = 0.541 for FST and 0.240 for TST; (**C**) Schematic representation of sociability and social novelty preference tests. ITI, intertrial interval. (**D** and **E**) Sociability tests. Both control and Ntf3-cKO mice spent significantly more time in investigating a mouse than an empty cage (**D**; n = 15 control mice and 12 Ntf3-cKO mice; Two-way ANOVA with post hoc Bonferroni's multiple comparisons: p=0.96 for genotype; p = 0.0306 for control group and 0.0346 for Ntf3-cKO group in social investigation time) but showed similar social preference index (**E**; Two-tailed unpaired t test, p = 0.781). (**F** and **G**) Social novelty preference tests. Both control and Ntf3-cKO mice spent significantly more time in investigating the stranger mouse than the familiar mouse (**F**; n = 15 control mice and 11 Ntf3-cKO mice; Two-way ANOVA with post hoc Bonferroni's multiple comparisons, p = 0.303 for genotype; p < 0.0001 for control group and p = 0.0011 for Ntf3-cKO group in social investigation time) but showed similar social discrimination index (**G**; Two-tailed unpaired t test, p = 0.94). Error bars represent SEM. ns = not significant, *p<0.05, **p<0.01, and ****p<0.0001.

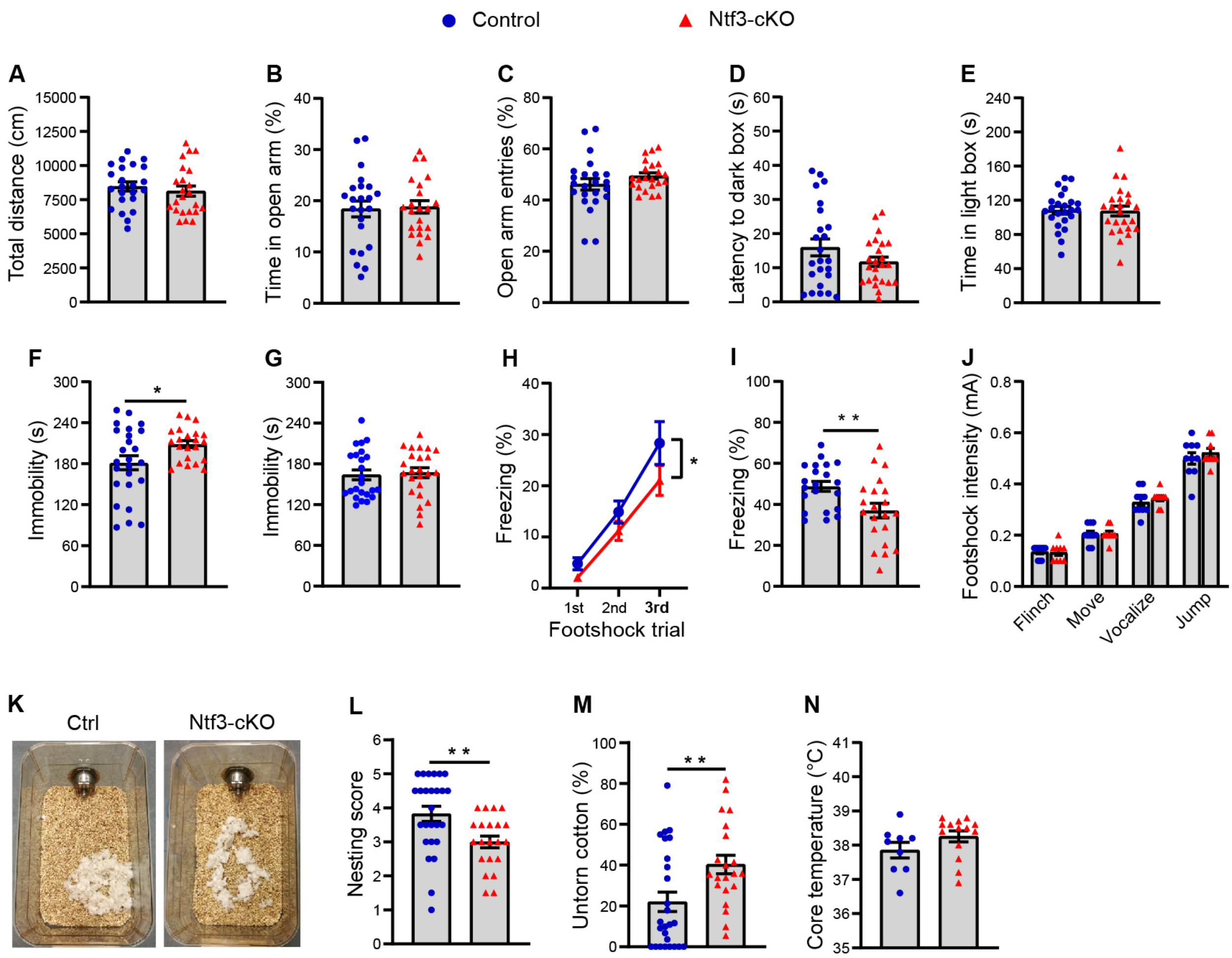


**Fig. S4 Female Ntf3-cKO mice displayed deficits in contextual fear memory and nest building.** (**A**) Total travel distance of mice in open field tests. n = 23 mice per genotype; Two-tailed unpaired t test, p = 0.491. (**B** and **C**) Time spent in the open arm and frequency of entry into the open arm in elevated plus-maze tests. n = 23 control mice and 22 Ntf3-cKO mice; Two-tailed unpaired t test, p = 0.835 for time in the open arm and p = 0.204 for open arm entries. (**D** and **E**) Latency to enter the dark box and time spent in the light box in light-dark box tests. n = 24 mice per genotype; Two-tailed unpaired t test, p = 0.144 for latency to enter the dark box and 0.907 for time spent in the light box. (**F**) Ntf3-cKO mice spent longer in immobility than control mice in FST. n = 25 control mice and 22 Ntf3-cKO mice; Two-tailed unpaired t test, p = 0.0247. (**G**) Control and Ntf3-cKO mice exhibited similar immobility in TST. n = 24 mice per genotype; Two-tailed unpaired t test, p = 0.745. (**H**) Ntf3-cKO mice showed fewer freezing responses to footshock than control mice during contextual fear conditioning. n = 19 control mice and 21 Ntf3-cKO mice; Two-way ANOVA, p < 0.0226 for genotype. (**I**) Ntf3-cKO mice spent less time in freezing than control mice during contextual fear memory retrieval. n = 19 control mice and 21 Ntf3-cKO mice; Two-tailed unpaired t test, p = 0.0088. (**J**) Control and Ntf3-cKO mice showed comparable pain threshold to footshock. n = 8 control mice and 9 Ntf3-cKO mice; Two-way ANOVA with post hoc Bonferroni's multiple comparisons, p = 0.377 for genotype. (**K**) Representative nests built by control and Ntf3-cKO mice. (**L**) Scores of nests built by control and Ntf3-cKO mice. n = 26 control mice and 21 Ntf3-cKO mice; Two-tailed Mann Whitney test, p<0.0037. (**M**) Percentage of untorn cotton after 12-hour nest building period. n = 26 control mice and 21 Ntf3-cKO mice; Two-tailed unpaired t test, p = 0.0089. (**N**) Control and Ntf3-cKO mice had similar core body temperature. n = 9 control mice and 14 Ntf3-cKO mice; Two-tailed unpaired t test, p = 0.155. Error bars represent SEM. *p<0.05 and **p<0.01.

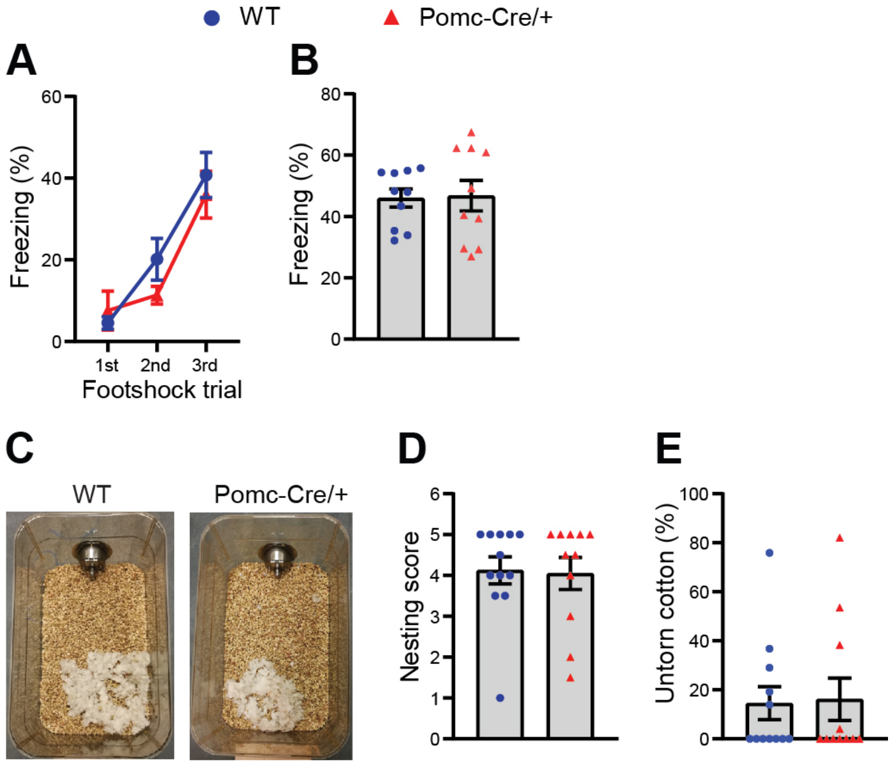


**Fig. S5 Male Pomc-Cre mice have normal contextual fear memory and nest building behavior.** (**A** and **B**) WT and Pomc-Cre/+ mice displayed comparable levels of freezing during contextual fear memory acquisition and retrieval. n = 10 mice per genotype. (**A**) Memory acquisition, two-way ANOVA with post hoc Bonferroni's multiple comparisons, p = 0.336 for genotype. (**B**) Memory retrieval, two-tailed unpaired t test, p = 0.899. (**C** to **E**) Nest building behavior. (**C**) Representative nests built by WT and Pomc-Cre/+ mice. (**D**) Scores of built nests. n = 12 WT mice and 11 Pomc-Cre/+ mice; Two-tailed Mann Whitney test, p = 0.913. (**E**) Percentage of untorn cotton. n = 12 WT mice and 11 Pomc-Cre/+ mice; Two-tailed unpaired t test, p = 0.884.

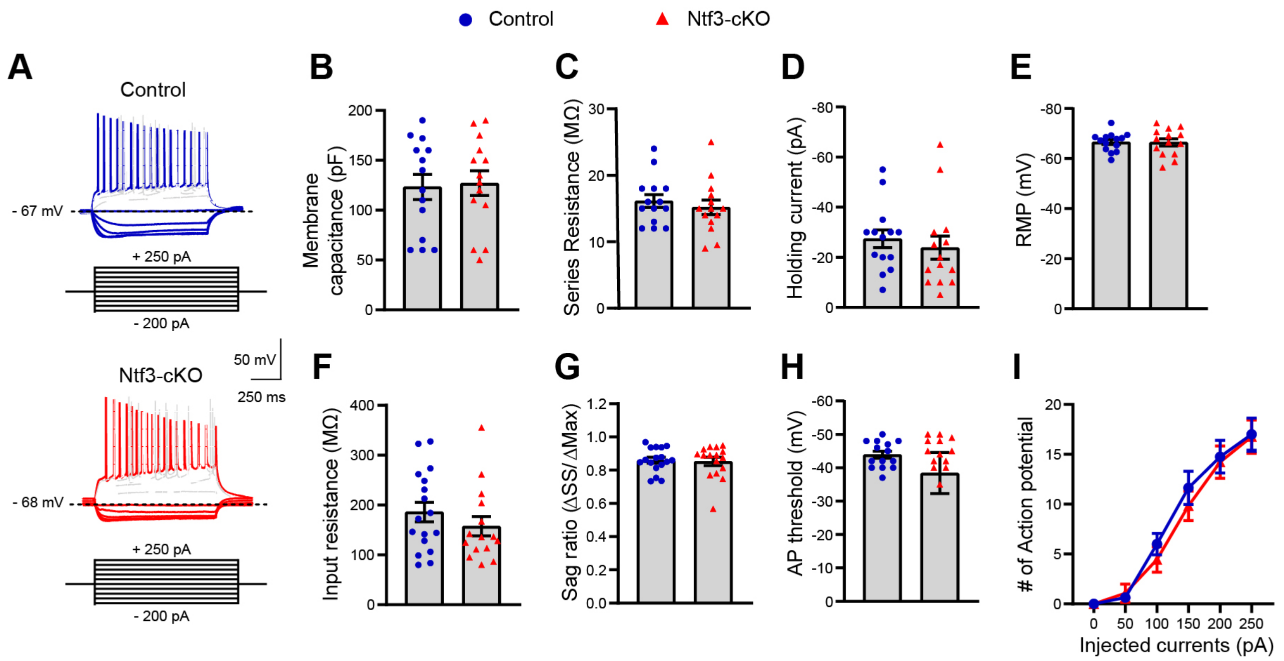


**Fig. S6 Normal intrinsic excitability of hippocampal CA3 pyramidal neurons in male Nft3-cKO mice.** (**A**) Representative traces of membrane potential responses to current injection. (**B**) Normal membrane capacitance in Ntf3-cKO neurons. n = 14 neurons from 7 control mice and 14 neurons from 4 Ntf3-cKO mice. Two-tailed unpaired t test, p = 0.832. (**C**) Normal series resistance in Ntf3-cKO neurons. n = 14 neurons from 7 control mice and 14 neurons from 4 Ntf3-cKO mice. Two-tailed unpaired t test, p = 0.517. (**D**) Normal holding current in Ntf3-cKO neurons. N = 14 neurons from 7 control mice and 14 neurons from 4 Ntf3-cKO mice. Two-tailed unpaired t test, p = 0.546. (**E**) Normal resting membrane potential in Ntf3-cKO neurons. n = 14 neurons from 7 control mice and 14 neurons from 4 Ntf3-cKO mice. Two-tailed unpaired t test, p = 0.941. (**F**) Normal input resistance in Ntf3-cKO neurons. n = 17 neurons from 7 control mice and 15 neurons from 4 Ntf3-cKO mice. Two-tailed unpaired t test, p = 0.31. (**G**) Normal voltage Sag ratio in Ntf3-cKO neurons. n = 17 neurons from 7 control mice and 16 neurons from 4 Ntf3-cKO mice. Two-tailed unpaired t test, p = 0.817. (**H**) Normal action potential threshold in Ntf3-cKO neurons. n = 14 neurons from 7 control mice and 14 neurons from 4 Ntf3-cKO mice. Two-tailed unpaired t test, p = 0.386. (**I**) Normal action potential responses to current injections in Ntf3-cKO neurons. n = 17 neurons from 7 control mice and 13 neurons from 4 Ntf3-cKO mice. Two-way ANOVA with post hoc Bonferroni's multiple comparisons, p = 0.413 for genotype.

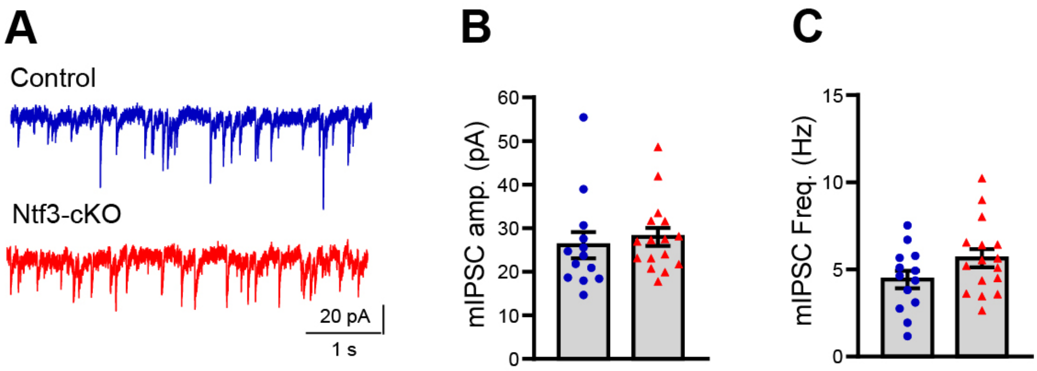


**Fig. S7 Deletion of the *Ntf3* gene in DG granule cells doesn't affect either amplitude or frequency of miniature inhibitory postsynaptic currents.** (**A**) Representative traces of mIPSCs. (**B**) Quantification of mIPSC amplitude. (**C**) Quantification of mIPSC frequency. n=13 cells from 3 Ctrl mice and 16 cells from 3 Ntf3-cKO mice. Two-tailed unpaired t test, p=0.594 for amplitude and p=0.111 for frequency.
